# MediSyn: uncertainty-aware visualization of multiple biomedical datasets to support drug treatment selection

**DOI:** 10.1101/165878

**Authors:** Chen He, Luana Micallef, Zia-ur-Rehman Tanoli, Samuel Kaski, Tero Aittokallio, Giulio Jacucci

**Affiliations:** Department of Computer Science, University of Helsinki, Gustaf Hällströmin katu 2b, 00560, Helsinki, Finland; Department of Computer Science, Aalto University, Konemiehentie 2, 02150 Espoo, Finland; Institute for Molecular Medicine Finland, University of Helsinki, 00014 Helsinki, Finland

**Keywords:** Interactive Visualization, Uncertainty Visualization, Multiple Datasets

## Abstract

**Background:** Dispersed biomedical databases limit user exploration to generate structured knowledge. *Linked Data* unifies data structures and makes the dispersed data easy to search across resources, but it lacks supporting human cognition to achieve insights. In addition, potential errors in the data are difficult to detect in their free formats. Devising a visualization that synthesizes multiple sources in such a way that links between data sources are transparent, and uncertainties, such as data conflicts, are salient is challenging.

**Results:** To investigate the requirements and challenges of uncertainty-aware visualizations of linked data, we developed MediSyn, a system that synthesizes medical datasets to support drug treatment selection. It uses a matrix-based layout to visually link drugs, targets (e.g., mutations), and tumor types. Data uncertainties are salient in MediSyn; for example, (i) missing data are exposed in the matrix view of drug-target relations; (ii) inconsistencies between datasets are shown via overlaid layers; and (iii) data credibility is conveyed through links to data provenance.

**Conclusions:** Through the synthesis of two manually curated datasets, cancer treatment biomarkers and drug-target bioactivities, a use case shows how MediSyn effectively supports the discovery of drug-repurposing opportunities. A study with six domain experts indicated that MediSyn benefited the drug selection and data inconsistency discovery. Though linked publication sources supported user exploration for further information, the causes of inconsistencies were not easy to find. Additionally, MediSyn could embrace more patient data to increase its informativeness. We derive design implications from the findings.

## Background

In biomedicine, the fruits of numerous biological assays and clinical studies are buried in various sources, such as publications and clinical reports, waiting to be translated into better treatments for patients [1, 2]. To accelerate such clinical practice and medical research, literature mining as well as crowdsourcing-based data-curation techniques are used to extract and collect useful biomedical information from the dispersed sources. Encouragingly, many curated databases provide open access, e.g., DrugBank [3] and clinicaltrials.gov [4], which inevitably benefits biomedical advances [5].

However, the isolated nature of biomedical databases still hinders the sharing and discovery of knowledge. To answer a biomedical question, scientists need to laboriously explore available sources via multiple and heterogeneous search services and then struggle to combine the selected information into a structured solution [6]. Due to the tediousness of the search process and the high cost of the cognitive load in matching sources [7], the abundant information sources available are often underexplored [6]. The ineffectiveness of translating datasets into useful insights calls attention to the essential issue of data integration.

*Linked Data*, as an effort to use the Semantic Web to interrelate data, encourages people to publish uniformly structured data, such as using the Resource Description Framework (RDF), so as to lower the barriers to connect data from different sources [8]. Some significant linked biomedical data projects include Bio2RDF [9] and Open PHACTS [10]. Nonetheless, the data published in Uniform Resource Identifiers (URIs) and RDF structures benefit the computer to interpret and correlate relevant information, but they do not facilitate human cognition to achieve insight. Hence, an interactive visualization tool that effectively synthesizes multiple biomedical datasets is required [1].

On the other hand, missing data and data errors in mined or curated biomedical datasets are difficult to detect in their free formats. Still, few efforts have been devoted to visualizing such data uncertainty to help biologists better understand the data [11]. Apart from that, the integration of multiple biomedical datasets brings another dimension of uncertainty: data consistency. Consistent information from different sources reinforces itself, giving people more confidence in the knowledge they acquire [12, 13], whereas conflicting data can motivate researchers to further explore the data sources to understand the causality. Visualization that conveys such uncertainty information among biomedical datasets [14] allows the user to make a more informed decision, such as treatment selection.

The purpose of this research was to visually synthesize multiple biomedical datasets, while exposing the uncertainties of the datasets to arouse user awareness of uncertain information and to facilitate drug treatment selection. In this paper, we present MediSyn (Fig. 1). It uses a matrix-based layout to correlate multiple drugs, targets, and tumor types. *Target* in this paper refers to mutations and wild-type genes. Sorting functions bring more relevant drugs to the front of the view to assist visual comparison of drug effects on multiple targets. The transparent representation and user exploration of drug-target relations enable the discovery of drug-repurposing opportunities, which is one contribution of this system.

**Figure 1.**
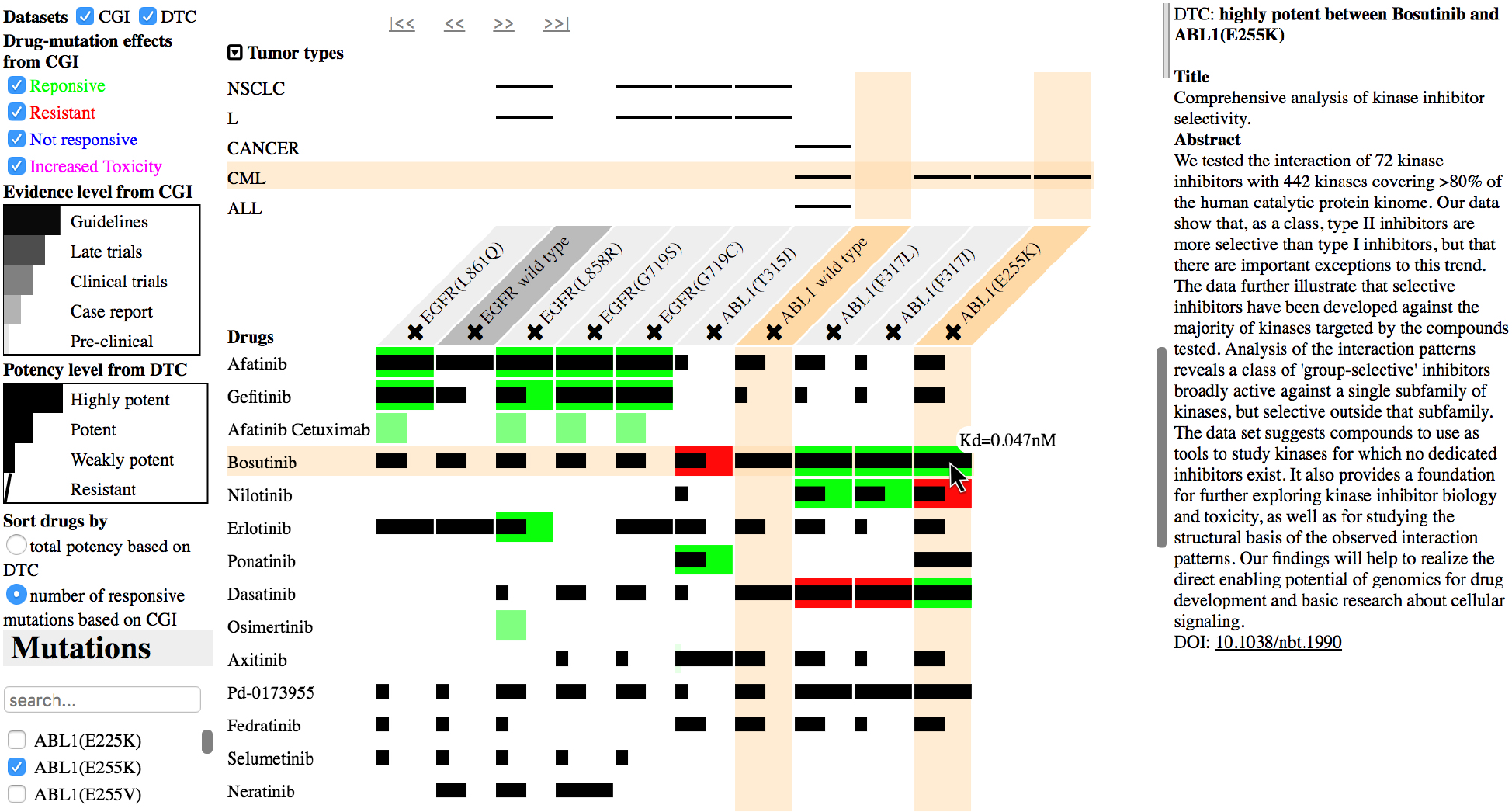
Overview of MediSyn. MediSyn synthesizes two biomedical datasets: cancer treatment biomarkers from Cancer Genome Interpreter (CGI) and drug-target bioactivities from Drug Target Commons (DTC). The left list contains the controllers, such as selection and sorting, whereas the middle view represents drug-target relations. Retrieved details of the clicked drug-target relation are shown on the right side. Here the user clicked on the potency bar of the drug bosutinib and mutation ABL1(E255K) so that the details, including the dataset, bioactivity state, and publication source, are shown on the right side.

Another contribution is that such a system visualizes data uncertainty to increase user awareness of data trustworthiness. First, the holistic relation representation among drugs, targets, and tumor types exposes missing data. Second, depicting datasets in overlaid layers enables the user to identify data consistency states from different sources. Third, visual encodings of different levels of clinical evidence expose data credibility. Data provenance, such as publications, can be interactively retrieved to convey the credibility of information sources.

MediSyn is implemented with two manually curated datasets, cancer treatment biomarkers from Cancer Genome Interpreter (CGI) and drug-target bioactivities from Drug Target Commons (DTC). A preliminary study with six domain experts showed that the synthesis of two datasets can increase user satisfaction and efficiency and lower choice difficulty in drug selection compared to user exploration with currently unlinked datasets. Subjective results showed positive feedback on MediSyn, such as simplicity and ease of use. Among others, the links to data sources, such as publications, appear to be an important and useful feature for the user to verify or acquire additional information about the data. The study results also indicated MediSyn effectively supported the discovery of data inconsistencies, but the causes of inconsistencies were not easy to find. Additionally, more patient data sources can be integrated to increase the informativeness of MediSyn.

Based on the results of the user study, we derive a set of design implications of MediSyn to inform two design problems: how to depict the correlated biomedical datasets; how to effectively expose and visually communicate data uncertainties.

## Related Work

To facilitate knowledge discovery from dispersed and heterogeneous biomedical datasets, some projects, such as Linked TCGA (The Cancer Genome Atlas Database) [15] and Open PHACTS [10], brought together pharmacological data resources and built data infrastructures to allow for the integration and interoperation of biomedical data. Several visualization tools have been built on top of the linked biomedical data platforms to support knowledge exploration, e.g., GenomeSnip [16] and PharmaTrek [17]. GenomeSnip [16], consisting of Genomic Wheel and Genomic Tracks, integrates knowledge of the human genome from multiple sources to support the exploration and cognition of the relationships between different genomic features. Genomic Wheel visualizes the hierarchical information of chromosomes, ideograms, genes, and cancer point mutations in circular layers, whereas Genomic Tracks visualizes gene information retrieved from Linked TCGA in tabular panels.

PharmaTrek [17] is based on Open PHACTS but integrates information on molecule-protein interactions and ligand structures to support multitarget drug discovery. It uses a heatmap to depict molecule and target activity values. The user can filter related molecules by setting the range for the activity values to each target, and he or she can retrieve additional targets related to the displayed molecules. In a similar two-dimensional layout, Campbell et al. [18] brought together biological, chemical, and clinical resources and built a confidence-based drug-target landscape along two evidence dimensions on a scatter plot. The x-axis of the scatter plot indicates ordered categories that provide evidence connecting proteins to disease, whereas the y-axis denotes ordered categories of evidence supporting small-molecule druggability for proteins.

These visualizations do not explicitly separate different datasets but rather take the linked data as a whole to facilitate user exploration across data sources. Similarly, some visual search platforms have been built in this manner to aid biomedical search across resources.

TripleMap [19] allows user exploration of biomedical entities, such as compounds, diseases, and assays [1]. It uses a node-link diagram to automatically connect user-selected entities based on the semantic tags retrieved from RDF datasets. ReVeaLD [20] has a visual query builder to help the user formulate a query in an intuitive way, and it displays results in a faceted results browser through a federated search.

Because trust in information requires an awareness of its provenance [21], we argue that users should be aware of information sources and have control of the sources, which can be based on their confidence in the datasets.

Several research efforts visualize datasets in separate views and then use linking and brushing techniques or explicit links to show data relations. ConTour [22] provides a relationship view of datasets, such as genes, compounds, and pathways, in columns at the bottom with a detailed view of the selected items above. The user selection of items in one column can highlight relevant items in other columns. Sorting and filtering functions can be flexibly combined to drill down into the data space. Similarly, StratomeX [23], based on VisBricks [24], employs a column-based layout to represent datasets, with bricks in those columns encoding potential subtypes or stratifications (partitionings into homogeneous subsets) of the data. Ribbons connect bricks of neighboring columns, with their width encoding the amount of data they share. Such explicit links are adopted in Domino [25] as well, which interrelates items between separate views of datasets using line connections. It enables the user to freely arrange and combine the blocks to tailor to the task at hand. For example, assembling Sankey diagrams [26] to recognize the flow in the datasets.

Different from the previous work, MediSyn uses overlaid layers to represent datasets not only to link but also to allow for comparison between datasets. Additionally, a matrix-based layout is adopted due to its scalability in visualizing data items as well as its support for the comparison of rows and columns. For instance, Bertifier [27] adopts a matrix-layout to link two data items, but cells visually encode a single data attribute associated with the item in that row and column. Lamy et al. proposed a matrix-based set visualization (rainbow boxes) with drugs in columns and their contraindications in rows to allow for the comparison of relevant drugs [28], but no indication of data sources is involved in the visualization.

Apart from that, MediSyn exploits the crucial but underexplored problem in biomedical data, that is, data uncertainty, to support a more-informed treatment selection.

## MediSyn

MediSyn (Fig. 1) is a matrix-based interactive visualization that supports drug treatment selection under uncertainty. It consists of three parts (Fig. 1). The left list contains the controllers, including dataset and mutation selection and sorting functions. The middle part is the matrix-based view with overlaid layers synthesizing datasets. The right part displays the detailed descriptions and the sources of user-clicked data.

### Datasets

Two manually curated datasets of drug-target relations are synthesized (Fig. 2). One is cancer treatment biomarkers from the Cancer Genome Interpreter (CGI) [29], and the other is drug-target bioactivities from Drug Target Commons (DTC) [30]. The CGI contains drug responses such as responsiveness and resistance to various mutations in different tumor types. Five *evidence levels*, i.e., pre-clinical, case report, early trials, late trials, and guidelines, such as Food and Drug Administration (FDA) guidelines – from the lowest to the highest – indicate the approval status of a drug. DTC contains bioactivities between different drugs and targets, which can be considered pre-clinical evidence. Due to the fact that the data from the CGI have a generally higher evidence level than those from DTC, we place higher priority on the data from the CGI in visual encodings.

**Figure 2.**
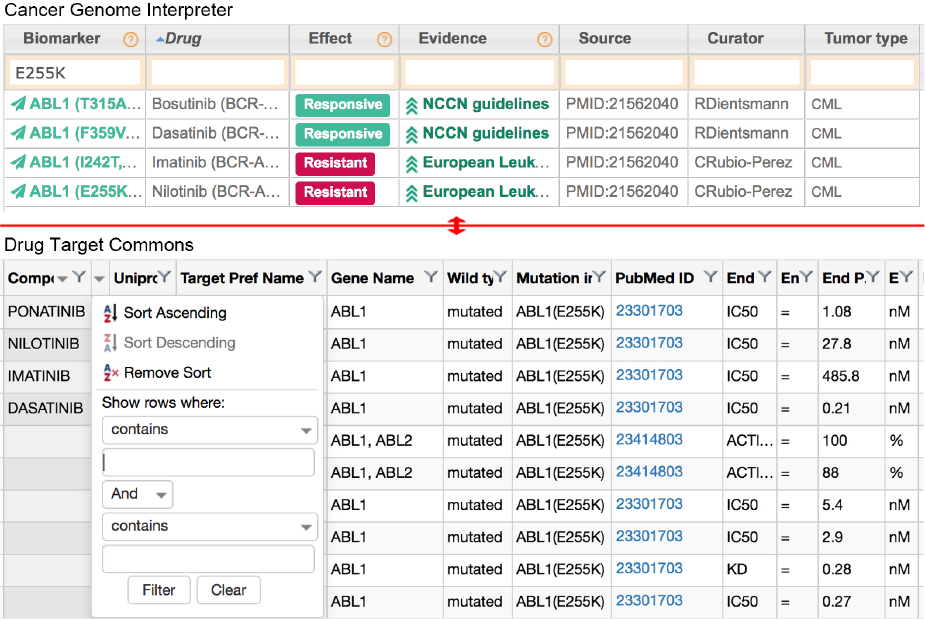
The upper part is a cancer treatment biomarker dataset from the CGI, whereas the lower part is a bioactivity dataset from DTC. We used the two webpages as the baseline interface in the user study.

Each bioactivity in DTC is described by a measurement type, such as Kd, Ki, and IC50, and the bioactivity value. We further categorize the bioactivity values to potency levels to make them easier for the user to understand. An activity value between 0 and 10 nM is classified as highly potent; a value between 10 and 1,000 nM denotes potency; a value between 1,000 and 10,000 nM indicates the drug is weakly potent; and a value over 10,000 nM indicates the drug is inactive [29]. If multiple bioactivities exist for the drug and target pair, we take the median as the activity value to avoid the disturbance of outliers.

### Visualization Design

The visualization supports a one-to-one representation of the relations between drugs, targets, and tumor types. It uses a matrix-based layout where each column represents a user-selected target. The rows above the targets represent tumor types, and the rows below depict related drugs (Fig. 3). Two overlaid layers representing the two datasets respectively visualize the relations between drugs and targets.

We prioritize the data parameters, abstract them to different data types, e.g., nominal and quantitative, and then map them to visual variables considering Mackinlay’s ranking [31], a ranking of visual variables regarding how accurately humans perform the corresponding perceptual task for different types of data. As depicted in Table 1, drugs and targets are nominal data with the highest priority. Thus, we encode them by position, which helps with forming the rows and columns of the matrix. Tumor types are nominal data and are related to targets only. We encode them by position as well, forming the rows at the top of the matrix (Fig. 3). If the mutation has been described in the tumor type, a horizontal line will appear in the corresponding table cell, which is inspired by linear diagrams representing relations of sets [32].

**Figure 3.**
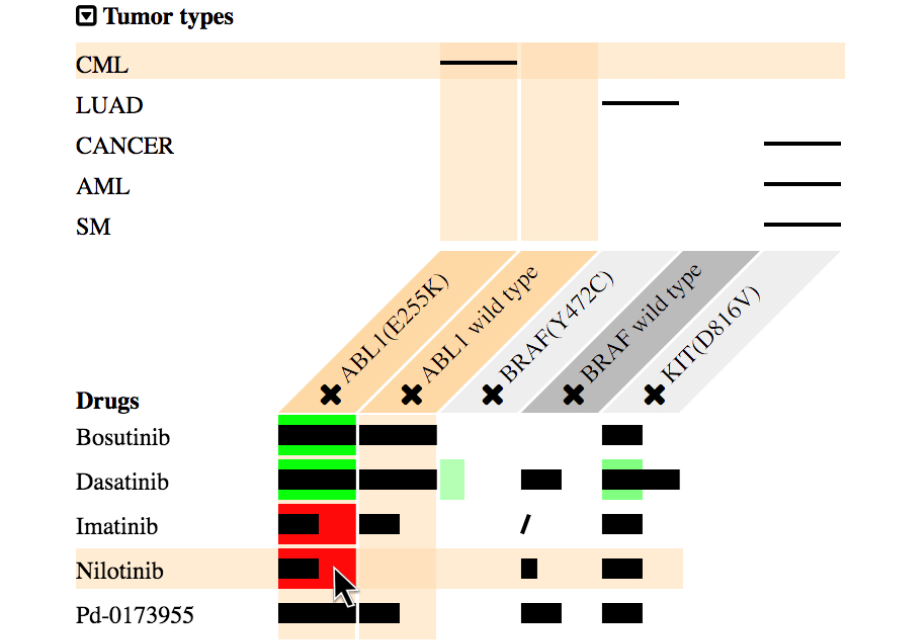
The matrix-based layout of MediSyn represents the relations among multiple drugs, targets, and tumor types. The overlaid layers depict the data from two datasets respectively with effcient visual encodings for prioritized data parameters.

**Table 1.** Visual variables encoding different parameters of our datasets based on their priority and importance in supporting drug treatment selection.

| Parameters | Data types | Visual variables |
| --- | --- | --- |
| Drug | Nominal | Position |
| Target | Nominal | Position |
| • Tumor type | Nominal | Position |
| Drug-target relation from CGI |  |  |
| Drug-target effect | Nominal | Hue |
| • Evidence level | Ordinal | Position, length, saturation |
| Drug-target relation from DTC |  |  |
| Bioactivity potency level | Ordinal | Position, length |

As we have fixed the positions of drug-target effects from the CGI in the corresponding cells, these nominal data then adopt the second-best visual variable, which is hue. Responsive effects are shown in green, whereas resistant effects are displayed in red (Fig. 3). The evidence levels of the biomarkers are ordinal data, which use a combination of position, length, and saturation encodings. As a result, the encodings of drug-target effects and their evidence levels translate the data into colored bars, i.e., each column of the matrix contains a vertically aligned bar chart. Finally, bioactivity potency levels from DTC are ordinal data residing in the corresponding cells as well, which constitute another layer of data on top of the CGI. We encode them by position and length. As illustrated in Fig. 3, the black bars on top of the colored bars with decreased width depict the potency levels of drug-target bioactivities, whereas slashes denote inactive bioactivities.

### Interaction Design

The interactions enable user exploration of the relations between multiple drugs and targets. The sorting functionalities based on different criteria support the user in identifying effective drugs Detailed descriptions as well as data provenance of the drug-target relations can be retrieved on demand.

#### Dataset and target selection

Based on the information from multiple sources, MediSyn allows the user to explore the relations of interested mutations to relevant drugs, tumor types, and the wild-type gene.

The user can choose to display the data from only interested or trusted data sources through controlling the checkboxes on the left top of Fig. 1.

Once a mutation is selected from the left list (Fig. 1), it is added as a new column in the matrix. All drugs related to the selected mutation are added as rows automatically. In addition, tumor types related to the selected mutation are retrieved and displayed above the matrix.

The wild-type gene of the selected mutation, if it exists in the datasets, is also added as a column. The wild type can be used to predict possible side effects of the drug. If the drug shows greater potency toward the wild type than the mutated gene, then possible side effects can be anticipated from this drug. A cross icon attached to the header of each column allows the user to remove the target.

#### Sorting

Sorting allows the user to rank the drugs based on different criteria to explore their relations to multiple targets. MediSyn allows the user to sort the drugs in three ways. If the user clicks the column header of a target, all drugs related to this target come to the top. The drugs containing data from both datasets come first; the drugs with data only from the CGI come second, whereas the ones described only in DTC come third, in descending order of the potency values, as the CGI data have a higher evidence level than the DTC data. Using the sort control on the left, the user can either sort the drugs by the sum of the potency values of all selected mutations to each drug based on DTC data or by the number of responsive mutations of each drug based on the CGI data. Both methods sort the drugs in descending order.

#### Highlighting

Highlighting provides visual cues to interrelate drugs, targets, and tumor types to the current focused data. Hovering over the drug name, i.e., row header, highlights all of its related targets as well as its related tumor types. Hovering over a bar highlights its mutation and drug as well as the column of its wild-type gene, if it exists. Hovering over the tumor name highlights all its related mutations.

#### Details on demand

Following Shneiderman’s information-seeking mantra [33], details regarding the drug-target relation as well as the data provenance are provided on demand. As the mouse hovers over the DTC bars, the detailed bioactivity values are shown as a tooltip. If the user clicks on any of the CGI or DTC bars, related information is shown on the right, including the dataset to which it belongs, a description of the drug-target relation, and the sources of the curated data, such as the title, abstract, and digital object identifier (DOI) of the publication. Clicking on the DOI of the publication will bring the user to the publication page.

### Visualizing Data Uncertainties

MediSyn uses a matrix-based layout coupled with overlaid layers to relate data items and synthesize datasets. Three types of data uncertainties are exposed in MediSyn to increase user awareness of data trustworthiness: missing data, data consistency, and data credibility. The matrix-based layout interrelates drugs, targets, and tumor types to facilitate user’s cognition of **missing data**.

A superimposed view facilitates direct comparison of data from multiple sources and exposes **data consistency** states. In our case, both datasets indicate drug-target relations, which allows them to share the same spatial mapping [34]. At the same time, direct comparison of data consistency is crucial in this case. Based on these two conditions, we adopted a superimposed view [34], i.e., overlaid data layers.

Overlaid layers of comparable data elements allow the user to easily perceive data consistency between datasets. Fig. 3 contains some inconsistent drug-target relations, where for the same drug and mutation pair the CGI value indicates resistance between them, whereas the DTC dataset shows the drug is potent toward the mutation, i.e., red bars overlaid with black potent bars. On the other hand, cases exist where the two datasets provide consistent results. For example, the cells where highly potent bars lie on top of the fully saturated green bars in Fig. 3.

**Data credibility** can be assessed in two ways. First, visual encodings of the evidence levels consisting of position, length, and saturation inform the credibility level of the drug-target relations from the CGI. Second, links to data sources, such as publications, can be retrieved on demand to expose data credibility.

### Implementation

MediSyn is implemented using D3.js [35]. It contains 536 different point mutations, among which 350 come from the CGI, 217 are from DTC, and 31 exist in both datasets (Table 2). Sixteen wild-type genes all come from DTC. There are in total 258 different kinds of drugs or drug combinations, 166 of which are from CGI, 116 are from DTC, and 24 exist in both datasets. A total of 2,405 different pairs of drug-target relations exist, 546 of which are cancer treatment biomarkers. The rest are from DTC, among which 665 pairs are wild-type drug interactions. Forty-two drug-mutation pairs contain data from both datasets. Finally, 52 tumor types are all retrieved from the CGI. MediSyn is available at http://medisyn.hiit.fi.

**Table 2.** Statistics of the data from the CGI and DTC.

|  | Mutation | Wild-type gene | Drug | Drug-target relation | Tumor type |
| --- | --- | --- | --- | --- | --- |
| CGI | 350 | 0 | 166 | 546 | 52 |
| DTC | 217 | 16 | 116 | 1,859 (665 wild-type targets) | 0 |
| Both | 31 | 0 | 24 | 42 | 0 |
| Total | 536 | 16 | 258 | 2,363 | 52 |

## Use Case

Studies have shown that even oncologists at a leading cancer center express low confidence in their knowledge of genomics [36]. MediSyn makes the knowledge of genomically informed therapy accessible and evaluable to clinicians. Such personalized cancer medicine involving the patient’s molecular profile, i.e., patient mutations, can be more advantageous than current standard therapies across tumor types [36].

As a use case, Fig. 4 shows the T315I mutation confers resistance to the majority of approved ABL1 inhibitors [37], such as the drug bosutinib, except for ponatinib, which has toxicity limitations. However, MediSyn exposes that axitinib could be a promising treatment for patients with the otherwise highly drug resistant mutation BCR-ABL1(T315I)-driven chronic myeloid leukemia, based on pre-clinical evidence from both datasets (highlighted drug in Fig. 4), which is also in agreement with the findings of Pemovska et al. [37]. This demonstrates how comprehensive representation of drug-target data can lead to unexpected and novel drug-repurposing opportunities.

**Figure 4.**
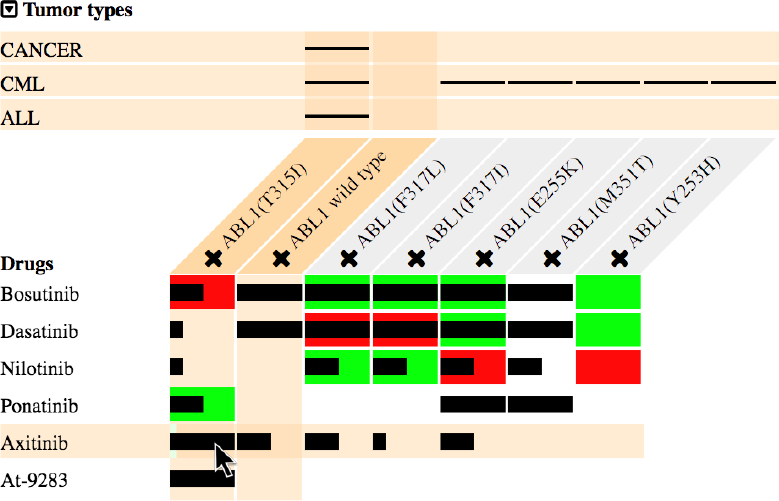
A case of the discovery of a drug-repurposing opportunity. Axitinib is promising to treat otherwise highly drug resistant mutation ABL1(T315I)-driven chronic myeloid leukemia based on pre-clinical evidence from both datasets.

In facilitating the identification of data uncertainty, the highlighted cell in Fig. 3, for instance, shows that according to the CGI, ABL1(E255K) is resistant to nilotinib, whereas DTC data show the drug is potent for this mutation. The user can find the same information from the original CGI and DTC webpages in Fig. 2, but such data conflicts are difficult to detect when the datasets are unlinked. In the user study, we inquired of a number of bioinformaticians about the possible cause of such inconsistencies. They provided some hypotheses but did not have an explicit answer (see the next section).

## User Study

To investigate the benefits of MediSyn as well as other possible insights and future design challenges resulting from data integration and uncertainty visualization, we did a within-participant study with six domain experts. To concretize the investigation, we raised the following two research questions. In addition, we assessed the user experience with the synthesized interface to select effective drugs.

- RQ1: What features of MediSyn are useful? What features need to be further developed?
- RQ2: Can MediSyn convey data inconsistencies to the user? How will user awareness of inconsistencies among datasets affect user trust in the curated data and in MediSyn?

We conducted the evaluation in a lab setting using the Chrome browser on a 13.3-inch MacBook Pro with a 2.8-GHz Intel Core i5 processor, 16 GB of RAM, and a built-in trackpad and keyboard. The display resolution was 2,560 * 1,600 pixels.

### Baseline

We used the original CGI and DTC webpages as the baseline system (Fig. 2) to assess the impact of MediSyn as a synthesized interface. The CGI cancer treatment biomarker page describes the mutations, drugs, evidence levels, data sources, and tumor types of the biomarkers in a table, as shown in the upper part of Fig. 2. The user can reorder the rows by clicking on the header of the column and can filter the rows using the filtering box at the top of each column. The DTC Web application allows the user to search bioactivities by a point mutation. It displays the relations of mutations, drugs, activity types, and values in a table as well. Similar to the CGI, the user can sort and filter the bioactivities using the control, as shown in the lower part of Fig. 2. For both datasets, clicking on the data source will open a new window that shows the source page of the curated data, such as a publication page.

### Participants

Among the six participants (three females; age mean: 28.6, SD: 5.32, N: 6), five were bioinformaticians, and one was a computer scientist. Participants were asked to complete a pre-questionnaire using a seven-point Likert scale so that their background and prior knowledge could be established (Table 3). Among the six participants, one participant claimed to use DTC occasionally but had never used the CGI. This participant was not quite familiar with the features of DTC (five on a seven-point Likert scale). Another participant stated that he had used the CGI before but not DTC and was not quite familiar with the features of the CGI (four on a seven-point Likert scale). The remaining four participants had never used either of the systems. The participants had little familiarity with cancer biomarkers (mean: 3.33, SD: 1.51, N: 6), drug-target bioactivities (mean: 2.83, SD: 1.83, N: 6), cancer drivers (mean: 2.83, SD: 1.94, N: 6), and anti-cancer drugs (mean: 3.17, SD: 1.72, N: 6). Also, they had no particular familiarity with tabular visualizations (mean: 4.83, SD: 2.14, N: 6). The participants thought that knowing the provenance of the displayed data was important (mean: 6.33, SD: 1.03, N: 6).

**Table 3.** Statistics on the prior knowledge of participants on a seven-point Likert scale.

|  | Mean | SD |
| --- | --- | --- |
| Familiarity with |  |  |
| Cancer treatment biomarkers | 3.33 | 1.51 |
| Drug-target bioactivities | 2.83 | 1.83 |
| Cancer drivers | 2.83 | 1.94 |
| Different kinds of anti-cancer drugs | 3.17 | 1.72 |
| Tabular visualization | 4.83 | 2.14 |
| Knowing the provenance of the displayed data is important. | 6.33 | 1.03 |

### Tasks

#### Task 1 (T1) - Drug selection

Each participant used both the baseline system (Fig. 2) and MediSyn (Fig. 1) to find the most effective drug for a pair of mutations. The order was counter-balanced. For each system, the participants used a different pair of mutations. We assigned two pairs of mutations. All four mutations had data in both datasets and had similar drug responsiveness data in the two datasets.

#### Task 2 (T2) - Inconsistency discovery

Each participant used MediSyn to find the inconsistency in the data between two datasets for a pair of mutations. We assigned a pair of mutations that contained inconsistency information from the datasets for both mutations.

### Procedure

Before T1 with each system, the participants were first trained on how to use the system. Training was active, as participants were asked to complete some basic tasks using the system through a printed introductory document. The experimenter ensured the participants understood how to complete these tasks before the actual experiment commenced. The whole training session took around 10 to 15 minutes for each system. During the actual tasks, participants were allowed to use pen and paper.

The participants then completed a questionnaire for each system in T1. The questionnaire adopted the design of ResQue [38] and Knijnenburg et al. [39]. For T2, participants were encouraged to think aloud while exploring the datasets. Afterward, the participants completed another questionnaire on the trustworthiness of the system as well as the curated data. Finally, we asked some interview questions to gather general feedback. The whole experiment took around an hour for each participant. Each participant was given a movie ticket as compensation. For each task, the screen was recorded and used for subsequent analysis.

### Overview of Results

#### Task performance

Fig. 5 shows the time spent with each system for T1 and the time spend on T2. For T1, using the baseline, the longest time spent was 12m20s (participant 5), and the shortest was 03m30s (participant 4). With MediSyn, the longest time was 03m50s (participant 5), and the shortest time was 01m10s (participant 6). On average, the six participants spent 6m22s with the baseline system (SD: 03m09s), and 1m57s when using MediSyn (SD: 01m00s). The participants required more than three times the time with the baseline than with MediSyn.

**Figure 5.**
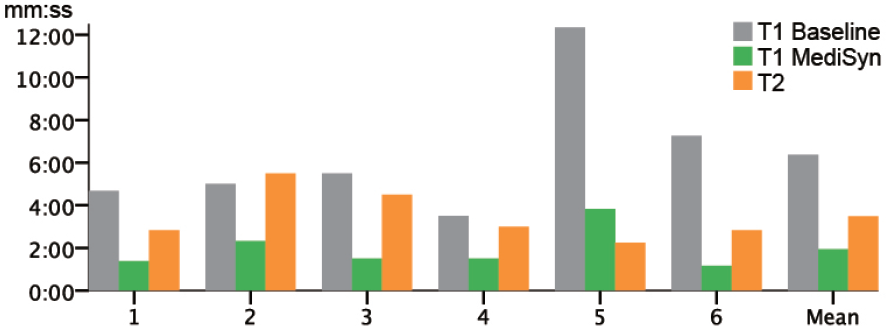
The time participants spent on drug selection and inconsistency discovery tasks. The x-axis indicates different participants and the aggregated mean, and the y-axis denotes the time in the format of mm:ss.

All participants eventually got the right answer for both systems during T1. The right answer was the drug that was responsive to both mutations based on CGI evidence and that had the lowest bioactivity value in DTC. Due to the small intersections of the two datasets, we could not set up a more complex task, such as drug selection for a group of four mutations.

For T2, the longest time spent was 05m30s (participant 2), whereas the shortest time was 02m15s (participant 5). On average, the participants spent 03m29s (SD: 01m14s). All participants found all the correct answers.

#### Questionnaire results

Fig. 6 shows user experience feedback for both systems. Participants were more satisfied with the selected drug using MediSyn (median: 5.5, N: 6) than with the baseline (median: 4, N: 6). They experienced less choice difficulty (MediSyn median: 2, baseline median: 3, N: 6). They also perceived MediSyn as easier to use (MediSyn median: 7, baseline median: 6, N: 6), which could also be observed in the improved task efficiency with MediSyn, and as requiring less effort (MediSyn median: 2, baseline median: 3, N: 6). These results can also be explained by the observation that the participants only needed the draft paper when working with the baseline. Participants tended to be more satisfied with and trusting in MediSyn (median: 6, N: 6) compared with the baseline (median: 5.5, N: 6). Similarly, they tended to use MediSyn again for drug selection tasks (MediSyn median: 6.5, baseline median: 6, N: 6). They tended to think that information provided in MediSyn (median: 5.5, N: 6) was more sufficient than the baseline (median: 5, N: 6).

**Figure 6.**
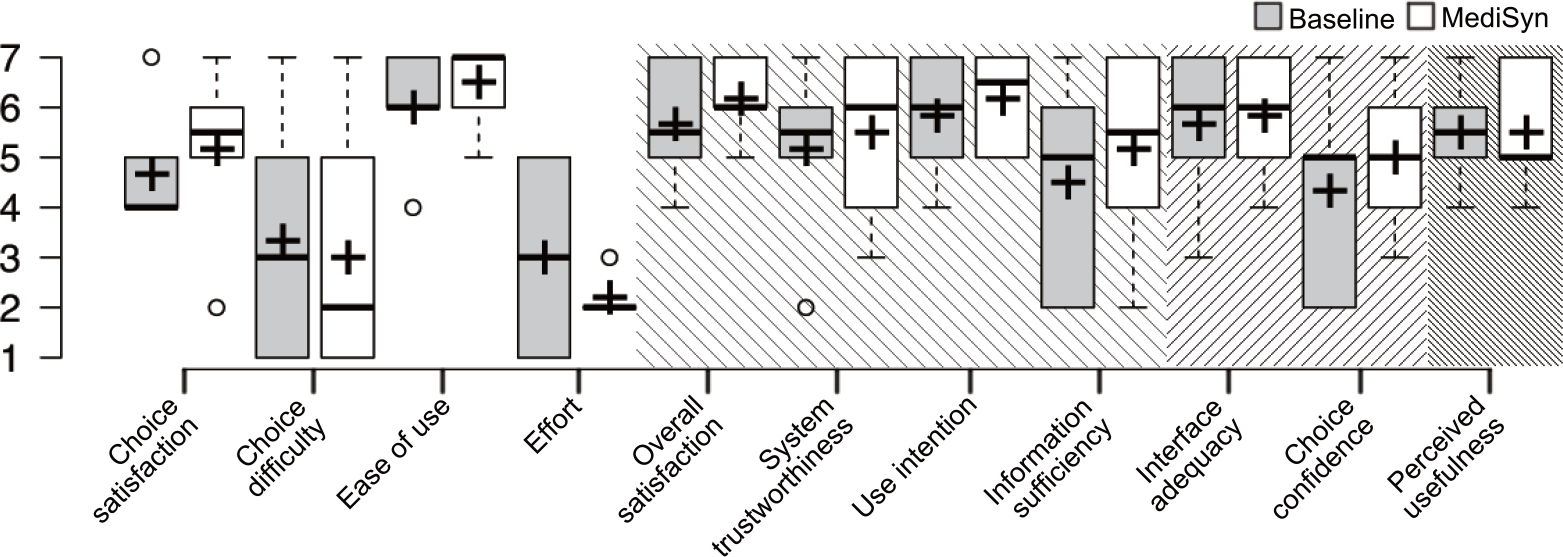
Results of the user experience with MediSyn and the baseline.

The results for interface adequacy (median: 6, N:6) and choice confidence (median: 5, N: 6) were the same for both systems, whereas MediSyn (median: 5, N: 6) was perceived as less useful compared with the baseline (median: 5.5, N: 6). A possible explanation could be that MediSyn extracted only some important data columns from DTC to display. Therefore, the users could find more abundant data properties for the bioactivities using the original DTC webpage.

Fig. 7 shows user trust feedback on MediSyn as well as on curated data. In general, the participants tended to think that manual data curation was error prone (the left most boxplot of Fig. 7, median: 3, N: 6). However, for these two manually curated datasets, they were unsure about the reliability of the data no matter whether they realized there existed inconsistencies in the datasets (median: 4, N: 6). On the other hand, before user perception of data inconsistency, i.e., during T1, the participants believed MediSyn was reliable (median: 6, N: 6). However, user trust in MediSyn tended to drop after participants found inconsistencies in the datasets during T2 (median: 5, N: 6).

**Figure 7.**
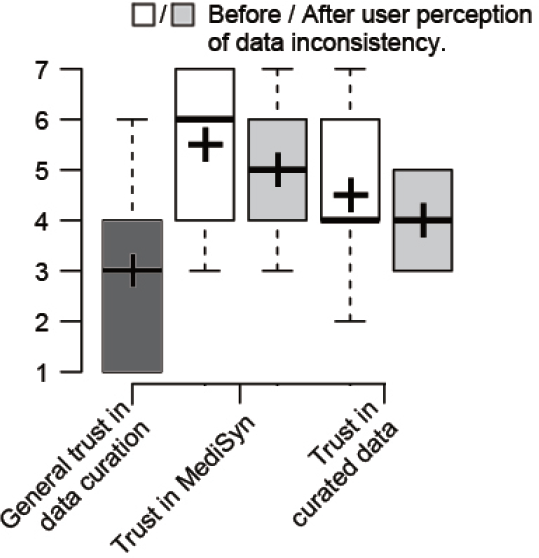
The results of general user trust in data curation as well as the user trust in the system and the curated data before / after the perception of data inconsistencies.

### Discussion and Design Implications

We discuss the results of the user study and derive a set of design implications to inform the design of future uncertainty-aware visual synthesizers for biomedical data.

#### The synthesis of datasets can increase choice satisfaction, lower choice difficulty, and improve task efficiency

Compared to unlinked datasets, the results of T1 showed the synthesized interface could improve efficiency and choice satisfaction and lower choice difficulty in drug selection. Two participants stated that MediSyn was simple and user friendly. Three participants suggested the visualization should have a better layout design; specifically, two participants said the information on the left was too dense, and one participant suggested stretching the bars because sometimes she could not tell if she was clicking on the CGI or DTC values. Two participants had difficulty matching the evidence levels to the legend. One participant could not understand the usage of different sorting functions.

#### The matrix view supports drug comparison and exposes missing data

The matrix-based view provides a scalable layout [40, 41] to support the perception of drug effects on multiple targets and tumor types, which enables the user to compare and select promising drugs for certain targets. Such a holistic view of drug-target relations also facilitates user cognition of **missing data**.

#### Depiction of datasets in overlaid layers facilitates direct comparison of data from multiple sources and effectively supports user perception of data consistency states

For T2, all participants found all conflicts for the designated mutations in a reasonable time frame. One participant expressed it was useful to have two datasets together, particularly for the second task. Otherwise, she could not realize there were inconsistencies in the datasets.

#### Exposed data inconsistencies tend to lower user trust in MediSyn but do not have observable effects of user trust in curated data

Most participants used the two datasets for the first time during the evaluation. They were unsure about the reliability of the datasets throughout the study. However, their trust in MediSyn tended to drop along with the cognitive transition from unawareness of the existence of data inconsistencies during T1 to the realization of their existence in T2.

#### No explicit answer was acquired on the rationale for conflicts in drug effects

Three participants stated that the inconsistency could be caused by patient complexity. For example, the patient could have acquired resistance due to a history of drug treatments. One participant declared the inconsistency could be due to the different measurements in experiments. For instance, in one case, the data from the CGI used the IC50 measurement type, whereas the DTC data used the Kd value. The rationale behind the data conflicts remains an open question, inviting the user to further investigate.

#### User accessibility to data sources of the curated data is an important and useful feature

Three participants expressed it was useful to have the link to publications easily accessible, which is in accordance with the pre-questionnaire result that shows knowing the provenance of the displayed data is important to the user. Two participants stated they would still need to read the paper before making the decision in T1. The tight coupling of data and the provenance allows the user to verify the credibility of the information and acquire additional information about the data.

#### More patient data need to be embraced to expand user knowledge

For T1, one participant asserted he would not decide on the treatment based only on patient mutations and needed to look for other information. Two participants claimed that patient data such as age and treatment history were also important to consider. One participant suggested taking a patient cell sample to experiment with the selected drug. How to embrace more information sources in an intuitive manner to further broaden the user’s knowledge of decision-making while avoiding the information overload problem remains a future research challenge.

#### Overview and details on demand support the scalability of the number of datasets

MediSyn displays datasets in overlaid layers. Based on the user study, such a superimposed view can effectively convey the states of data consistency. However, it can also cause visual clutter with the increase of the number of datasets [34]. In practice, if we have more than two datasets, we propose using MediSyn to provide an overview of the data from available sources. For instance, each data cell in MediSyn can depict the possibility of resistance between the drug and target as well as that of responsiveness based on the calculation across all sources. The user can have control over the weight of the data sources in the calculation. With this informative overview, the user can then further explore the details of the data cells.

## Conclusions

In this paper, we presented MediSyn, an uncertainty aware interactive visualization that synthesizes biomedical datasets to support drug treatment selection. The matrix view coupled with overlaid layers presents a comprehensive relation among drugs, targets, and tumor types from multiple sources, supporting the comparison of drug effects on multiple targets. A use case with the implementation of MediSyn synthesizing two datasets, cancer treatment biomarkers from the CGI and drug-target bioactivities from DTC, showed its effectiveness in supporting the discovery of drug repurposing opportunities.

From a visualization research perspective, MediSyn visualizes the uncertainty of the datasets to support more informed decision making. The matrix-based layout exposes missing data. Overlaid layers ease the perception of data consistency. Visual encodings of evidence levels as well as links to data provenance convey data credibility.

A preliminary study with six domain experts showed that such a synthesized interface can increase choice satisfaction and efficiency and lower choice difficulty compared to currently unlinked datasets in supporting drug selection. Subjective results showed generally positive feedback. User accessibility to data sources, among other factors, appears to be a crucial and useful feature. Additionally, MediSyn facilitates user perception of data inconsistencies, but the cause of conflicts remains an open question.

MediSyn is still in its early stage and has great potential to be improved. First, the layout and readability of the visual design can be improved to ease the perception of the links between the datasets and data properties. Second, the drugs can be linked to diseases to further benefit the discovery of drug-repurposing opportunities. Third, enabling the user to sort the columns based on the activities of mutations can further refine the user selection of drug treatment. Because not all driver genes are equally important in the course of tumorigenesis [42]. Tumors may be more addicted to mutations in certain drivers, which provide basic capabilities to cancer cells [42]. Fourth, we plan to incorporate more information sources, one of which is the clinicaltrials.gov dataset containing basic patient information of drug clinical tests, to further enhance user knowledge. Fifth, design implications of MediSyn can be generalized to serve other types of data collections.

## List of abbreviations used

RDF: Resource Description Framework; URI: Uniform Resource Identifiers; TCGA: The Cancer Genome Atlas Database; Open PHACTS: Open Pharmacological Concept Triple Store; CGI: Cancer Genome Interpreter; DTC: Drug Target Commons.

## Declarations

Ethics approval and consent to participate

We recruited the participants for our study on a voluntary basis. All participants were rewarded with a movie ticket for their voluntary participation. All participants read and signed a consent form regarding their voluntary participation, their right to withdrawal, the data logging and confidentiality before the actual study.

## Consent for publication

Not applicable.

## Availability of data and materials

The datasets used by MediSyn are downloadable at https://www.cancergenomeinterpreter.org/biomarkers and https://drugtargetcommons.fimm.fi/. Data collected in the user study cannot be disclosed to third parties, as stated in the study consent form.

## Authors’ contributions

SK, TA, and GJ conceived the idea behind this work. LM and CH designed the visualization techniques and user study. CH implemented MediSyn, conducted the study, analyzed the collected data, and drafted the manuscript. ZT provided the DTC dataset. ZT, TA, and GJ provided useful suggestions during the iterative revision of MediSyn. TA proposed the representative use case of MediSyn. GJ and LM provided critical revisions to the writing. All authors read and approved the final manuscript.

## Acknowledgements

We would like to thank Baris Serim from Department of Computer Science and Zaid Alam from Institute for Molecular Medicine Finland, University of Helsinki for their valuable discussions and insightful suggestions during the design of MediSyn.

